# The development of visual acuity and crowding reveals the slow fine-tuning of foveal vision

**DOI:** 10.1101/2025.03.27.645750

**Authors:** John A. Greenwood, Marilia Kyprianou, Tessa M. Dekker

## Abstract

The adult visual system is characterised by high-resolution foveal vision and a peripheral field limited by crowding, the disruption to object recognition in clutter that gives a summary ‘gist’ over fine detail. In children, crowding is elevated foveally, with the estimated age where foveal crowding drops to adult-like levels varying widely from 5 to 12+ years. As crowding restricts key processes like reading, characterisation of this developmental trajectory is critical. Using methods optimised to measure crowding in children, adults and typically-developing children (n=119; 3-13 years) judged the orientation of a foveal ‘VacMan’ target either in isolation or surrounded by ‘ghost’ flankers. For isolated elements, acuity (measured as gap-size thresholds) dropped rapidly to adult-like levels at 5-6 years. Thresholds rose when flanked/crowded, with elevations highest at 3-4 years, persisting at 5-6 years, and dropping to adult-like levels at 7-8 years. A meta-analysis of our results and 13 prior studies reveals a consistent developmental trajectory, despite wide methodological variations. We further demonstrate that developmental crowding shows the same selectivity for target-flanker similarity as peripheral crowding, consistent with common mechanisms. This prolonged development reveals a shifting balance in the visual system between the processing of fine detail vs. the ‘gist’ of the scene.

## Introduction

The human visual system is *foveated* – our ability to recognise objects is greater in central (foveal) vision than in the periphery. Although the decline in peripheral vision is in part related to reduced acuity^1^, a greater disruption arises through *crowding*, a process whereby surrounding clutter interferes with the recognition of an otherwise visible target object^2,3^. This cortical process^4^ has been argued to provide a summary or ‘gist’ of the peripheral field (e.g. the predominant colour, orientation, etc. in a given region), at the expense of fine detail^5^. In the fovea, where our sensitivity to fine detail is greatest, crowding is typically minimal^6^. This balance between fine detail and ‘gist’ differs in children, however, with pronounced elevations in crowding in their foveal vision^7^. These prolonged elevations are likely to limit key abilities like reading^8^ and visual search^9^, as well as making the visual system vulnerable to disruption, as seen with the further elevations in crowding in developmental disorders ranging from amblyopia^10^ to dyslexia^11^. Estimates of the age at which crowding becomes adult-like vary widely, however. Though acuity is adult-like at 5-6 years^12–14^, estimates of the age of maturity for crowding vary from 5 through to 12 years or later^15–23^. Our aim was to better characterise the developmental trajectory of crowding using procedures optimised to measure crowding in a child-friendly format, and to compare these measures of crowding with those from prior studies to gain a broader view of this developmental process.

Crowding disrupts the recognition of visual modalities ranging from orientation^24^ to colour and motion^25,26^, as well as complex elements such as faces^27,28^. In peripheral vision, crowding occurs when flanker objects fall within an interference zone around a target^29^. Interference zones increase in size with eccentricity^30,31^ such that the centre-to-centre separation of objects needs to be beyond around half the target eccentricity to avoid crowding, regardless of object size^31,32^. In the fovea, crowding effects are typically small, to the extent that early studies argued they were indistinguishable from processes such as masking^33,34^. Subsequent work has nonetheless demonstrated small but consistent foveal crowding effects^6,35^. The mechanisms underlying crowding are contested, with high-level accounts proposing that crowding derives from attentional resolution^36^ or Gestalt processes of grouping^37^. These high-level accounts cannot explain the systematic nature of crowded errors however, whereby errors made regarding a given target are not random but rather follow the appearance of the surrounding flankers^38^. Pooling models explain these errors as the unwanted combination of target and flanker elements^39^, with population pooling models^40^ in particular able to predict the errors made regarding foveal stimuli presented to children^41^. It is this pooling that provides the ‘gist’ of the visual scene, which may be an efficient way to represent information-rich scenes with limited resources^5,42^. In children, the elevations in foveal crowding suggest that this drive for pooling and the computation of the ‘gist’ of the scene occurs more extensively than in adults.

Numerous studies have demonstrated that foveal crowding is elevated in children relative to adults^15–23^, with both a higher level of disruption from flankers and an increased spatial extent over which flankers interfere with target recognition. Estimates for the age at which these elevations recede to adult-like levels vary widely, however, as shown in Figure 1. The earliest of these estimates suggest that the spatial extent of foveal crowding declines with age to reach adult-like levels around the age of 5-7 years^15,16^, obtained using target letters surrounded by flanker letters. Others using letters report a slightly later estimate of 7-9 years^21^, aligning with letter-chart based measurements suggesting maturity around 7 years^22^. Later estimates include measurements with Landolt-C and tumbling-E elements that show maturity at 9 years^18,19^, while others find crowding that is still immature at 11 years, suggesting maturity at 12 or beyond^20^. In addition to the above-noted stimulus differences, prior studies vary widely in their stimulus arrangement, with variations in the separation between elements particularly likely to impact the measures. Several studies varied the spacing between target and flankers directly, and quantified performance via edge-to-edge separation^18–20^, which may be suboptimal given that crowding is likely driven by centre-to-centre separation^31,32^ (though this has been contested in foveal vision^6,33^). Others have scaled the size and separation of elements^15,21,22,43^, though with widely-spaced elements that are less effective at estimating the strength of crowding than arrays with closely separated elements^44–46^. To resolve these conflicting conclusions about the development of crowding, we need to re-examine this developmental trajectory using approaches optimised for the measurement of crowding in both children and adults.

**Figure 1.**
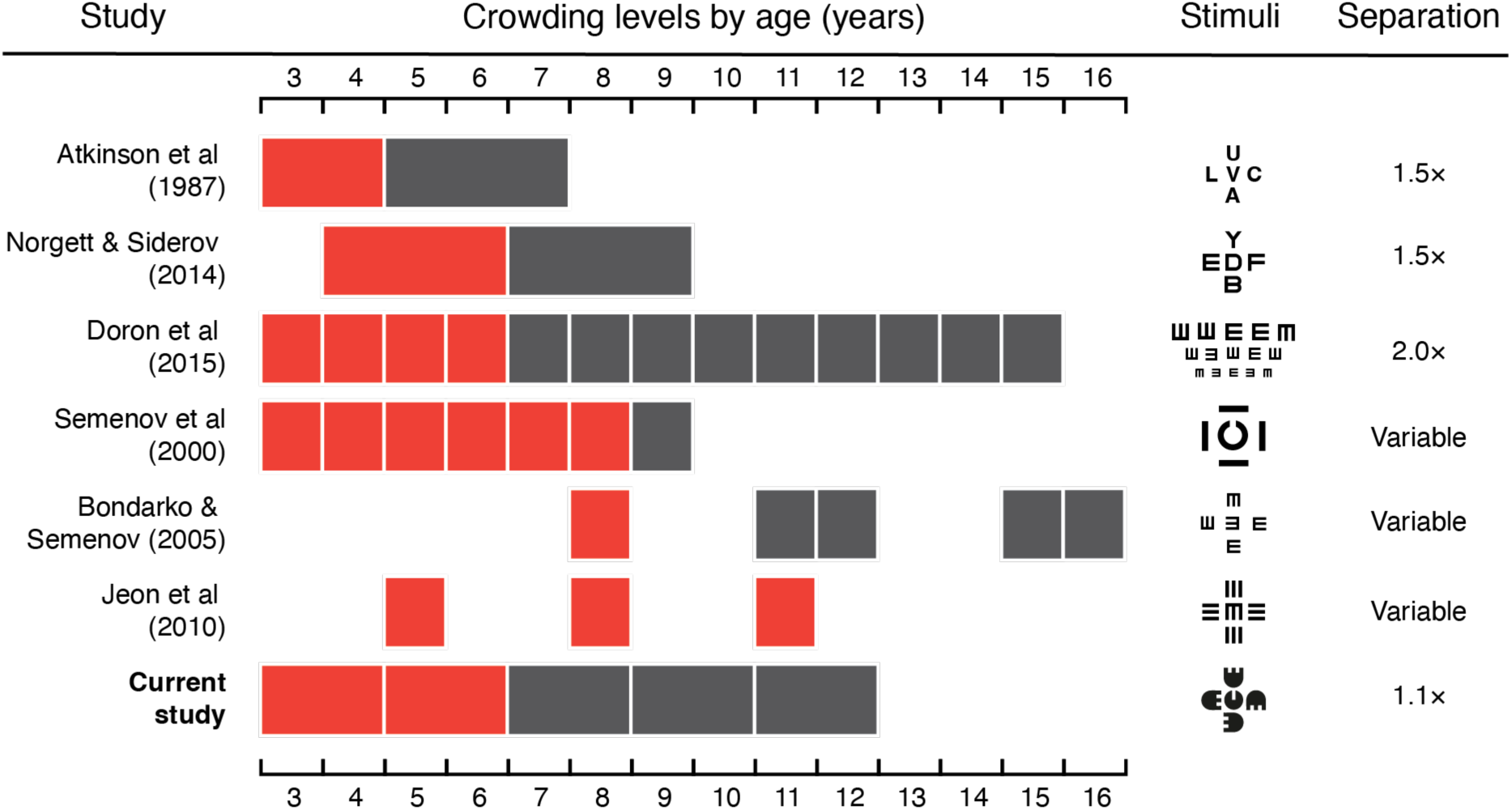
A comparison of estimates of the developmental trajectory of crowding, including the current study. We include only published studies where childhood crowding is compared directly against adult or late-adolescent crowding. Where multiple conditions were tested, we include those with the most ‘standard’ configuration (e.g. a target and 4 adjacent flankers). Red blocks show ages at which crowding levels were significantly elevated above those of adult participants; grey blocks show non-significant differences (‘adult-like’ performance). Block widths indicate the grouping of participants by age. Example stimuli are depicted for each study, with their centre-to-centre separation in multiples of element diameter, except where this property was directly manipulated (marked as ‘variable’). Studies are sorted by the age at which crowding becomes adult-like, excluding the current study.

With this aim, we developed child-friendly procedures adapted from those previously shown to optimally measure crowding in adult peripheral vision. Participants judged the orientation of a videogame character (Visual Acuity Man, or ‘VacMan’) viewed foveally, either in isolation or surrounded by ‘ghost’ flankers, as used previously in both typically developing children and those with amblyopia^41,47^. The advantage of this paradigm is its use of target and flanker elements that are similar enough to induce crowding, but with distinct identities introduced through the videogame-like context in order to enhance task comprehension and minimise source confusion. Given the optimality of measuring the spatial extent of crowding as the centre-to-centre separation between closely spaced elements^44^, we measured crowding using scaled elements with a centre-to-centre separation between the target and its flankers of 1.1x the stimulus size. We assessed acuity and crowding using independent measurements, with sizes varied using QUEST and catch trials included to ensure task engagement. A further advantage of this approach is that threshold estimates can be converted to reveal the spatial extent of crowding, as well as other metrics used to assess crowding previously, which we used to perform a meta-analysis of prior studies.

As well as characterising the developmental trajectory, these methods further allow us to address the debate on whether foveal and peripheral crowding derive from the same mechanism^6,31–34^. Although the scale of foveal crowding has been noted to make it difficult to distinguish from masking effects in the adult fovea^33,34^, the elevated crowding levels in children allow us to more clearly address this question. A key property of crowding in this regard is its selectivity for target-flanker similarity^48^ – crowding is reduced when target and flanker elements differ in properties such as contrast polarity, colour and depth. For instance, a black target amongst white flankers gives less crowding than an all-black array^48–50^. In typical adults, these effects have similarly been observed in the fovea^34,51^. In children, though there is some suggestion that varying flanker colour may reduce foveal crowding (with red flankers around a black target)^52^, it is unclear whether target-flanker differences reduce these disruptive effects as a general principle (i.e. whether they occur across other dimensions), as in adults. To fill this gap, we examined the selectivity of crowding for target-flanker similarity to contrast polarity in a subset of children, as well as the developmental trajectory of this selectivity.

We thus sought to examine both the age at which foveal crowding recedes to adult-like levels, and whether foveal crowding shows the same selectivity for target-flanker similarity as observed in peripheral vision. To this end, crowding and its selectivity was measured in children between the ages of 3-13, with comparison to adults conducting the same procedures.

## Methods

### Design

Children and adults judged the orientation of a black target element, known as Visual Acuity Man (‘VacMan’), either in isolation or flanked by four black ‘ghost’ elements (Figure 2). All participants completed these two conditions, with a subset also completing a third condition where the black target was surrounded by white flankers. Thresholds were measured in each condition to find the smallest gap size in the target element that allowed accurate recognition. When present, flankers were scaled in both size and separation, as described below. Ethical approval for all procedures was obtained from the UCL Experimental Psychology Research Ethics Committee (Project ID EP/11153/001), in accordance with the Declaration of Helsinki.

**Figure 2.**
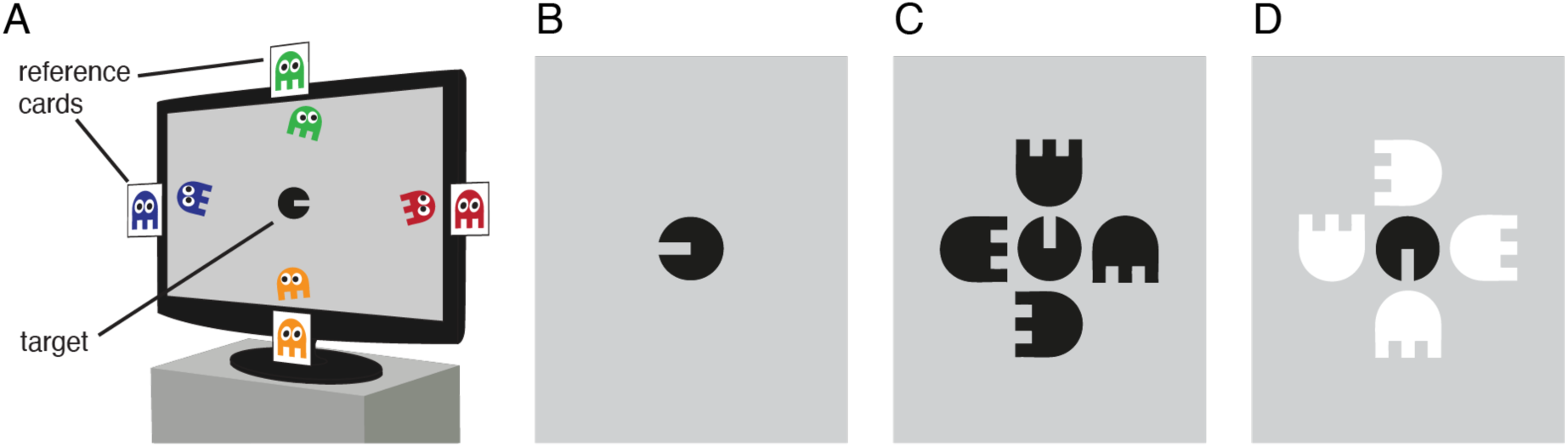
Example stimuli. **A**. Schematic depiction of the monitor display during unflanked trials, with Visual Acuity Man (VacMan) at the centre and four coloured reference ghosts moving around the monitor edge and presented on external cards to the side. **B**. An example target stimulus during unflanked trials. **C**. Example stimuli from the flanked-same condition, in which four same-polarity (black) flankers surrounded the target at random orientations. **D**. An example trial from the different-polarity condition, where flankers were white.

### Participants

A total of 125 participants were recruited, including 95 children and 30 adults. All were required to have normal or corrected-to-normal vision, with normal colour-naming abilities (checked using the stimuli prior to participation), and no history of neurological abnormalities. Participants were required to complete at least one block of trials to be included in the study. 3 children failed to meet these criteria, and a further 3 began testing but failed to complete the first condition. All 6 were thus excluded.

The final sample comprised 89 children (49 female) with ages between 3.2-13.1 years (mean: 8.2 years), and 30 adults (20 female, and including the 3 authors), with ages between 18.3-37.6 (mean: 23.1 years). For analysis, children were divided into 5 groups by age, with 10 3-4 year-olds (mean age 3.8 years, range 3.2-4.4), 22 5-6 year-olds (mean age 6.2, range 5.1-6.9), 24 7-8 year-olds (mean age 7.8, range 7.0-8.9), 14 9-10 year-olds (mean age 9.9, range 9.0-10.9 years), and 19 11-12 year-olds (mean age 12.2, range 11.1-13.1 years). Adults were paid for their participation and children rewarded with toys. Prior to participation, adult participants and the parents/carers of children gave informed consent, with children giving informed assent.

### Apparatus

Stimuli and experimental procedures were programmed using MATLAB (The MathWorks, Ltd., Cambridge, UK) on a Dell PC (Dell, Round Rock, TX) running PsychToolbox^53,54^. Stimuli were presented on a 27” ASUS VG278HE LCD monitor, with 1920 × 1080 resolution and 120 Hz refresh rate. The monitor was calibrated using a Minolta photometer (Konica Minolta Sensing Americas, Ramsey, NJ) and linearised in software, to give a maximum luminance of 222 cd/m^2^. Stimuli were viewed binocularly from 3 m, with the experimenter recording participants’ verbal responses via keypad.

### Stimuli and procedures

The experiment involved five video-game characters, as used previously^47^. The VacMan target was presented at the centre of the screen as a circle with a horizontal gap on one side, similar to a ‘filled-in’ Landolt-C (Figure 2B). The gap portrayed its ‘mouth’ and had a stroke width equivalent to one-fifth the stimulus diameter, as with Sloan letters^55^. During trials, the VacMan target was presented dark against the mid-gray background at a Weber contrast of 75%. Four achromatic ghost characters were also presented, with a semi-circular head and square ‘legs’ with gaps that were also one-fifth the stimulus diameter. When achromatic, the ghosts served as flanker stimuli in the flanked conditions (Figure 2C-D). In the unflanked condition, distinctly coloured animations of the ghosts (green above, red to the right, orange below, and blue to the left) moved along the screen boundaries (Figure 2A). Coloured ghosts were also shown on reference cards adhered to the edges of the monitor to serve as response options for the identification of the VacMan target orientation.

All participants began with the unflanked condition, where on each trial they reported which of the four ghosts VacMan was facing in order to help VacMan catch the ghost that he wanted to eat (a single-interval stimulus with a 4-Alternative Forced Choice response). Each ghost had a distinct colour and moved slowly along the monitor edges with a centre-to-centre separation approximately 5.5° from the target horizontally and 3.0° vertically (Figure 2A), making the chance of any crowding with the target unlikely. Responses based on ghost colour were used to avoid left-right response confusions^56^, though children could indicate the spatial location of the ghost if they wished. The use of clearly individuated characters also helped children with task comprehension, particularly in the flanked conditions – referring to the VacMan character helped make clear which element they were required to respond to, minimising the possibility of errors arising from source confusion^57^. Prior to testing, subjects were familiarised with the characters via printed cards illustrating VacMan, the ghosts and an example of a trial, through which normal colour-naming abilities were also checked. Feedback was given after each trial via brief animations whereby a colour-rendered VacMan either smiled or frowned. After every third correct response, a longer animation occurred where VacMan ran towards and “ate” the correct ghost. This feedback animation acted both as a reward and as a reminder of the task.

In the two flanked conditions, Vac-Man and the four ghost flankers were achromatic and presented without eyes to ensure that the target and its flankers were sufficiently similar to induce crowding. On each trial of these flanked conditions, the achromatic ghost-flankers were presented at a random orientation (with one at each cardinal orientation). Children were instructed that the ghosts became colourless to ‘hide’ from VacMan, but it was emphasised that they maintained the same colour/position mapping as before (e.g., as in the unflanked condition, the green ghost was above VacMan, the red to the right, etc.), which was reinforced through the presence of the reference cards. As before, participants reported the ghost that VacMan was facing by referring either to its colour (via the four reference cards at the monitor boundaries) or location. All children completed one flanked condition where target and flanker elements had the same contrast polarity (‘flanked same polarity’), with all elements presented dark against the mid-gray background at 75% Weber contrast (Figure 2C). A subset of children were also tested with a second flanked condition where a dark target (as in the other conditions) was flanked by light (‘white’) ghost-flankers, again at 75% Weber contrast (Figure 2D). 40 children and 10 adults completed this third condition.

In all three conditions, the size of the ghosts and VacMan (including the visibility of its “mouth”) was varied using QUEST^58^, set to converge on a gap size (for VacMan’s mouth gap) that gave 62.5% correct performance. Each block of trials consisted of a single QUEST staircase run on one of the conditions for 35 trials (not including practice trials). The target size was always scaled such that the diameter of VacMan was 5 times the mouth size. When flanked, the size of all elements was matched, with a centre-to-centre distance between the target and flankers set to 1.1 times the stimulus diameter. This separation is recommended by prior research as optimal for the robust measurement of crowding^44^, and has been used to measure crowding in both adults and children previously^41,59^. Scaling stimuli in this way allows measurement of both the magnitude of crowding (as the difference in thresholds between unflanked and flanked conditions) and its spatial extent (since the stimulus scaling allows conversion from gap-size thresholds to the centre-to-centre separation between elements, as we explore later).

The QUEST procedure was modified in four ways to increase its suitability for testing children. First, five practise trials were presented to begin each block of trials, where the gap size was 5.3’, well above any thresholds observed during pilot testing. These trials were not included in final analyses. If performance on these trials was poor, the task was explained again and the test re-started. Second, easier “catch” trials were presented periodically (on trials 5, 15, 25 and 35) where stimulus size was scaled to triple the current QUEST estimate of threshold. These catch trials were included both to minimise the frustration and loss of motivation that can occur when placing continuous trials near threshold, and to measure inattentiveness during the block, as recommended by prior work^60^. Third, to further reduce the effect of inattentiveness on threshold estimates, thresholds were determined by fitting psychometric functions to the data (as recommended previously^60^). Catch trials were included in the final data used in this fitting procedure. The fourth modification was related to this later fitting – because QUEST ordinarily places stimulus sizes close to threshold, the resulting range of values can be insufficient for curve fitting. Stimulus size on each trial was thus jittered using Gaussian noise added to the current QUEST threshold estimate (with a standard deviation of half the threshold estimate). This jitter increased the range of stimulus sizes, in turn improving the subsequent fitting of psychometric functions to the data.

We aimed for each condition to be completed two times by children and three for adults, though testing was stopped after at least one block in each condition if cooperation waned or the participant did not wish to continue. As above, all children completed the unflanked and flanked-same polarity conditions, with a subset also completing the third flanked-opposite polarity condition. In each block, participants would earn points after correct responses to each of the four catch trials. When children achieved high scores in each condition, they received stickers to exchange for a toy at the end of the experiment as a means to maintain motivation throughout the study. Rest breaks were permitted between the conditions. The full duration of the experiment was 30-60 minutes per person, followed by a debrief.

## Results

### Screening and analyses

The total number of trials completed per condition ranged from 35-105 (i.e. 1-3 blocks of trials). When split into the 5 age groups, this gave an average of 45.5 trials per condition for 3-4 year olds, 59.8 for 5-6 year olds, 51.1 for 7-8 year olds, 56.6 for 9-10 year olds, 51.5 for 11-12 year olds and 65.8 for the adult group.

Performance on catch trials with large stimulus sizes was uniformly high, as shown by age in Figure 3A. Each age group had some individuals who failed on at least one of the catch trials, which was slightly higher in the youngest 3-4 year old group. Mean percent-correct performance for both the unflanked and flanked same-polarity conditions was nonetheless above 98% in all groups and conditions save for the flanked condition in the 3-4 year olds, which dropped slightly to average 93.8% correct. Given that catch trials were correctly reported on the majority of trials in all cases, all of these children were included in the final analyses. Supplementary analyses were also run after removing children who scored below 100% correct on catch trials, which did not differ from the main analyses (Figure S1).

**Figure 3.**
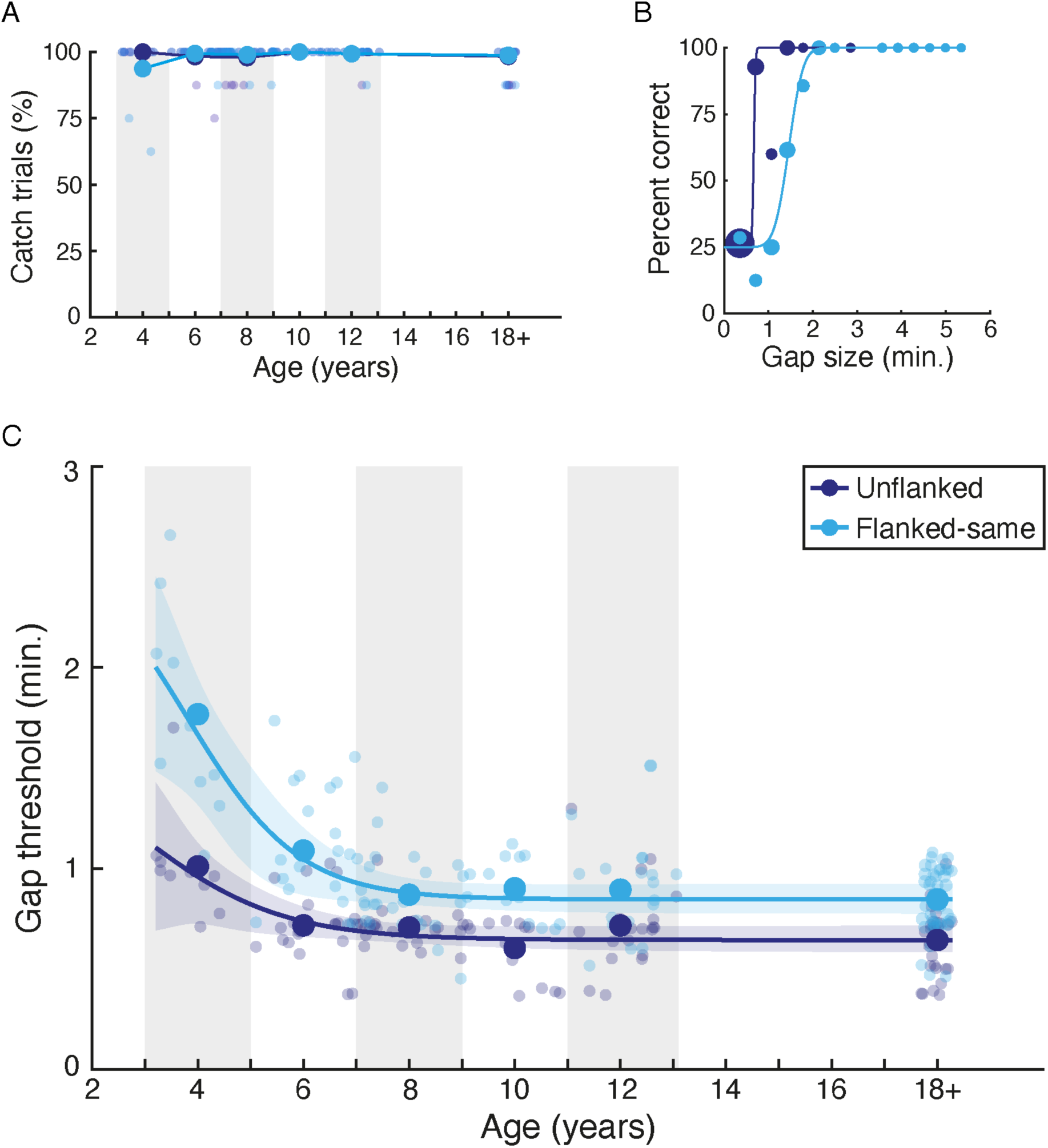
The development of acuity and crowding. **A.** Percent-correct performance on catch trials (large stimulus sizes) as a function of age. Individuals are shown as small points and the mean within each age range (shown via grey/white bands) as large points, separately for the unflanked (dark blue) and flanked-same (light blue) conditions. **B.** Example percent-correct data and best-fitting psychometric functions for one observer (age 6), plotted as a function of stimulus size separated for the unflanked (dark blue) and flanked-same (light blue) conditions. Dot size indicates the number of trials at each point. **C.** Gap-size thresholds in minutes of arc, plotted against age. Individuals are shown as small points and the mean in each age range as large points, separately for the unflanked and flanked-same conditions. Adult data has been collapsed to fall near 18 years. Lines show the best fitting logistic function to each dataset, with shaded regions showing the 95% range of the fits to bootstrapped data (1000 samples).

To determine the threshold gap size required for performance in each condition, repeated blocks of trials were first combined for each participant and stimulus condition, with the corresponding proportion correct scores collated for each stimulus size (including catch trials). Psychometric functions were then fit to the data using a cumulative Gaussian function with two free parameters (midpoint and slope). Given the variable number of trials at each gap size (driven by the variability in the QUEST algorithm, as above), these fits were determined by weighting the least-squared error by the number of trials at each point and seeking to minimise this value.

Example proportion correct values and best-fitting psychometric functions for one observer (aged 6) are shown in Figure 3B. The number of trials at each gap size is depicted via the size of each point, with the weighted psychometric function fit emphasising those with the most trials. Performance in the unflanked condition (dark blue) begins at chance (25% correct) for the smallest gaps and rises rapidly to ceiling as stimulus size increases. A slower progression from chance to ceiling is evident in the flanked-same condition (light blue) where the black target was surrounded by black flankers. For each condition, gap-size thresholds were taken from these psychometric functions at the size where performance reached 62.5% correct (midway between chance and ceiling).

### The developmental trajectory of acuity and crowding

Figure 3C shows the final gap-size thresholds in minutes of arc, plotted as a function of age. These values represent the smallest gap that can be identified in the VacMan target (at 62.5% correct) when unflanked (dark blue) or surrounded by flankers with the same polarity (light blue). A two-way mixed effects ANOVA was run to examine the overall pattern, with a between-subjects factor for age and a within-subjects factor for flanker condition. This analysis revealed a significant main effect of age (F_5,113_ = 20.85, p < .0001), reflecting the drop in thresholds with age, and a significant main effect of flanker condition (F_1,113_ = 193.50, p < .0001), reflecting the rise in thresholds when flankers were added. The interaction between age and flanker condition was also significant (F_5,113_ = 9.71, p < .0001), reflecting the larger differences between unflanked and flanked performance in the youngest children compared to the older cohorts.

To examine the effect of age in each condition, a series of planned contrasts were performed. Unflanked thresholds, a measure of visual acuity, were highest in the youngest children, but rapidly dropped to adult-like levels. This pattern is supported by independent samples t-tests between children in each age group and adults, which differed significantly with a very large effect size at 3-4 years (t_38_= 5.148, p < 0.0001, d = 1.88), but not at 5-6 years (t_50_ = 1.556, p= 0.126, d = 0.44) or beyond at 7-8 years (t_52_ = 1.643, p= 0.106, d = 0.45), 9-10 years (t_42_ = -0.755, p= 0.455, d = 0.24) or 11-12 years (t_47_ = 1.307, p= 0.198, d = 0.38). Acuity is thus elevated in young children aged 3-4 years and converges to adult levels at the age of 5-6 years.

Flanked thresholds were higher than unflanked thresholds for all age groups, following the main effect of crowding condition. To examine the development of crowding, flanked thresholds in each age group for children were compared with those of adults. Flanked thresholds were considerably elevated in 3-4 year olds relative to adults (t_38_ = 8.610, p < 0.0001, d = 3.14), with continued elevation at 5-6 years (t_50_ = 3.592, p= 0.001, d = 1.01), both with large-to-very-large effect sizes. Thresholds dropped to become indistinguishable from adults at 7-8 years, with a non-significant difference (t_52_ = 0.452, p = 0.653, d = 0.12), which remained flat thereafter with non-significant differences at 9-10 (t_42_ = 1.007, p= 0.320, d = 0.33) and 11-12 years (t_47_ = 0.812, p= 0.421, d = 0.24). Flanked performance thus indicates that crowding levels are initially strongly elevated in young children between the ages of 3-6, becoming indistinguishable from adult levels at the age of 7-8 years, later than the point at which acuity becomes adult-like.

As outlined in the introduction and Figure 1, prior studies of developmental crowding show a striking variability in the age at which crowding has been argued to become adult-like, ranging from 5 to 12+ years. Our estimate of 7-8 years for the point of maturity falls between these extremes. We next sought to investigate the cause of these discrepancies, both to build consensus around the developmental trajectory and to trace these differences back to their most likely methodological explanation (in turn considering the best-practice approach to measure crowding in children). With this aim, we first examined whether these discrepancies can be explained by different approaches to the measurement of crowding (specifically by varying flanker target-separation directly, or by scaling element size and separation), followed by a meta-analysis to gain a broader view of the developmental trajectory of crowding.

### A meta-analysis of the developmental trajectory of crowding

As outlined above, crowding has been measured using a variety of stimuli, procedures, and measurement approaches. One apparent division in Figure 1 is between studies where elements were scaled in both size and inter-element separation (as in the current study), which show earlier ages of maturation than those that varied separation directly using fixed-size elements^18–20^. Because the VacMan stimuli of the current study have been used to measure crowding previously using both the scaling approach (in the current study and previously^41^) and with direct manipulations of inter-element separation^47^, we can directly test whether thresholds differ under these two approaches. To do so, thresholds in all three studies were analysed using the same procedures (see Supplementary Information) and converted to give estimates of the centre-to-centre separation between elements at threshold. Thresholds for the centre-to-centre separation between elements in the flanked-same condition in these three studies are plotted in Figure S2. These analyses show a highly similar developmental trajectory in these three studies, suggesting that thresholds obtained using a scaled approach (as in the current study and previously^41^) are comparable to those that vary inter-element separation directly^47^.

We next sought a more direct comparison of our data against prior examinations of the developmental trajectory of crowding, both to obtain an overall consensus trajectory and to consider the methodological characteristics that can bias the estimates of individual studies away from the central tendency. A major complication that arises in comparing prior estimates is the wide range of metrics used to measure crowding. However, an advantage of the scaling approach in the present study (measuring flanked and unflanked thresholds by scaling element size and inter-element separation proportionally) is that it allows for conversion into the same units as these previously-used metrics. We thus undertook a series of comparisons between prior datasets and our threshold values after conversion to the same metric, building towards a meta-analysis of the developmental trajectory of crowding.

Methods for data extraction and conversion for each study are described in the Supplementary Information, along with individual comparisons between our values and those from 13 prior studies in their respective units of measurement (Figure S3). This includes both the studies shown in Figure 115,^18–22^ as well as others including unpublished datasets and those lacking direct comparison to adult performance levels^17,23,41,43,45,47,61^. Taking these analyses a step further, we then normalised each of these 13 datasets to span the same range, using our dataset range as a reference. The resulting estimates of normalised crowding across development are plotted in Figure 4.

**Figure 4.**
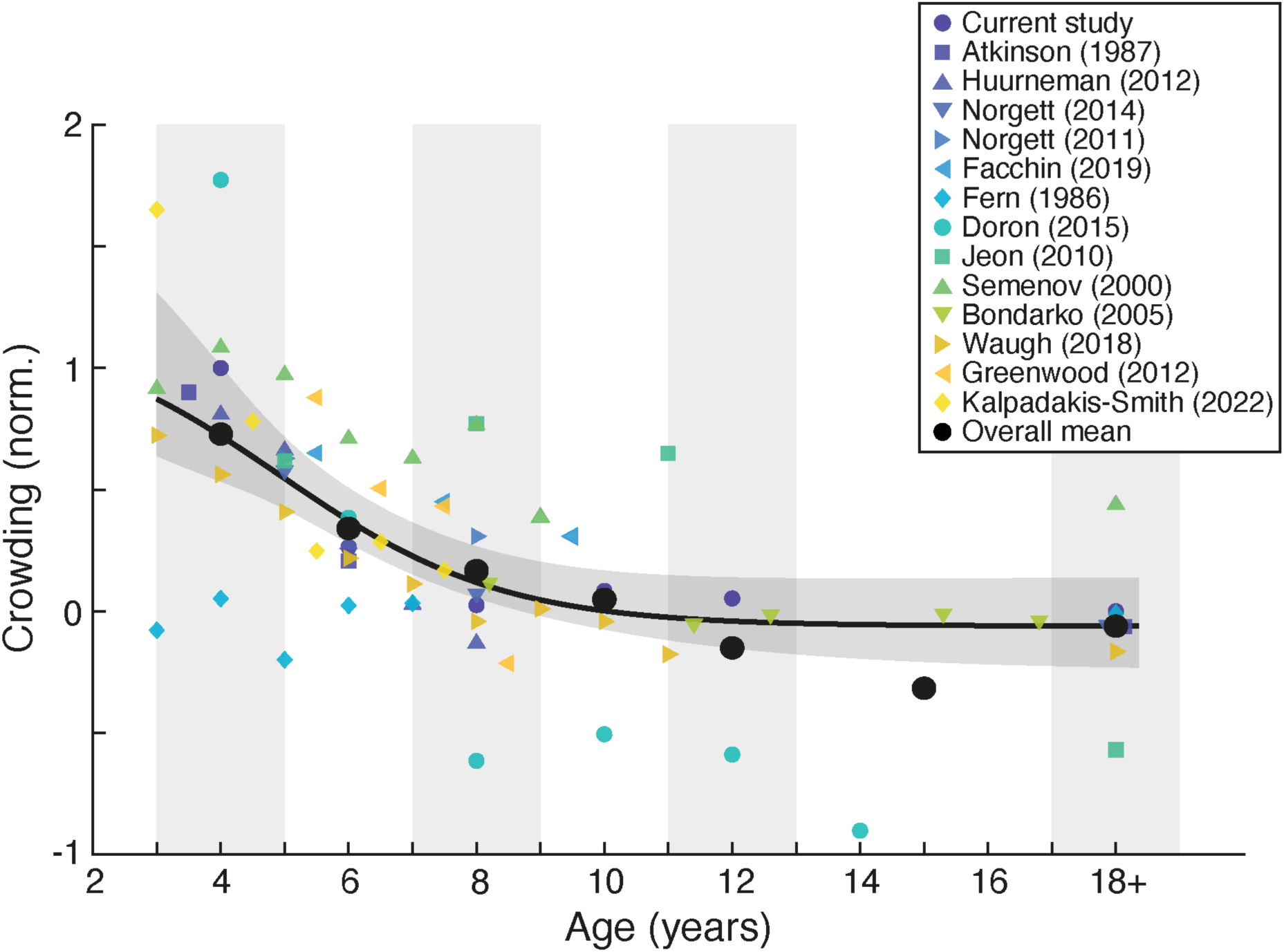
A meta-analysis of studies measuring the developmental trajectory of crowding. Crowding is plotted in normalised units for direct comparison (see Supplementary Information), as a function of age on the x-axis. Small symbols show estimates from individual studies (see legend), while the large black points show the mean within each age range (indicated via shaded regions). The black line is the best-fitting logistic function fit to all studies, with shaded region indicating the 95% range of the fits to 1000 bootstrapped samples. Some adult thresholds have been displaced on the x-axis for visibility.

Normalised crowding levels show some variation across studies, but an overall developmental trajectory is nonetheless apparent. To examine this trajectory statistically, values were pooled in similar age ranges to those of the current dataset. Crowding levels were highest in the 3-4 year age range, which differed significantly from adult levels (t_24_= 3.933, p < 0.001, d = 1.74). These levels remained significantly elevated at 5-6 years with a large effect size (t_21_ = 3.255, p= 0.004, d = 1.48), but dropped to levels that did not differ significantly from adults at 7-8 years (t_20_ = 1.436, p= 0.167, d = 0.66), and remained equivalent at 9-10 years (t_11_ = 0.571, p= 0.580, d = 0.32), 11-12 years (t_9_ = -0.482, p= 0.641, d = 0.30) and 13-17 years (t_8_ = -1.026, p= 0.335, d = 0.71). Although these analyses include repeated contributions from individual studies within a given age group, analyses conducted by binning individual datasets to give a single value for each age range produce the same pattern of results (Figure S4). This overall pattern thus follows the same trajectory as that of the current dataset, despite variations in stimulus type (including letters, symbols, and numbers), inter-element separation (from intermediate to narrow spacings, and studies that manipulate this property directly), and measurement approach (including the use of scaling vs. separation manipulations, the number of response options, use of single targets vs. letter-chart configurations, and so on).

Some divergence from this developmental trend is nonetheless apparent in individual studies, which can be traced to methodological differences. Although their estimated age of maturity agrees with ours, the estimates of Doron *et al*^22^ run higher than others in the youngest children. It is likely that this elevation reflects the use of high numbers of elements in a letter chart arrangement, which may cause uncertainty regarding the target response in the youngest children. Interestingly, these values then drop to very low levels in the oldest children (producing the slight dip in mean normalised crowding levels in the oldest groups, due to the small number of studies with data in this range). This too may reflect the letter-chart arrangement – the wide spacing would mean that each element is only flanked closely by up to 2 elements (depending on the letter position in each line), which would underestimate the low crowding levels at older ages in particular. Low estimates are also evident in the data of Fern et al^17^, who used widely spaced flanker bars surrounding target letters. The combination of wide spacing and the low similarity between target letters and flanker bars (as noted previously^21,44–46^) may have driven this underestimation.

Divergences are also apparent in the estimates of Jeon *et al* ^20^, who directly varied the edge-to-edge separation between target and flanker elements of a fixed size. These estimates are high and unvarying in children, but drop considerably in adults, where estimates are much lower than those of the current study and others reporting adult values. It may be that the adults tested by Jeon *et al*^20^ were more experienced with psychological testing than other samples, or that differences were present in testing procedures between children and adults that could underestimate crowding in the latter. We also show in the comparison to individual studies (Figure S3) that the use of relative measures (which combine estimates of acuity and crowding to express crowding as a multiple of acuity levels) lead to noisier estimates compared to studies where acuity and crowding are measured independently (as with the scaling approach of the current study). Nonetheless, Figure 4 shows that the majority of these 13 studies show good agreement with the current study, with a clear overall trend in the combined developmental trajectory, and an estimate of the age of maturity for crowding (7-8 years) matching that seen in Figure 3.

### The release from crowding

Given suggestions that foveal crowding may involve distinct processes in children and adults, we next sought to examine a key feature of crowding in peripheral vision – the selectivity of foveal crowding for target-flanker similarity. Differences in the contrast polarity of target and flanker elements have reliably been found to reduce the strength of crowding in peripheral vision^48–50^, with some suggestion that differences in target-flanker colour may also reduce foveal crowding in children^52^. Here we compare the foveal vision of children and adults to test whether this same selectivity is present for differences in contrast polarity, using a subset of children and adults who completed an additional condition where the black target was flanked by white flankers (‘flanked different’).

Gap-size thresholds from 40 children and 10 adults are presented in Figure 5. Thresholds in the unflanked condition follow the same developmental trajectory as shown in Figure 3, with elevations in the youngest children that quickly recede to adult-like levels. Performance in the flanked-same condition is also similar to that of Figure 3, with elevations evident at all ages, but largest in the youngest children. Thresholds in the flanked-different condition were reduced relative to the flanked-same condition, though did not reach the levels of unflanked performance. Given the reduced sample size, statistical tests were run by taking a median split of the children by age, giving one group of 3-6 year olds and another of 7-12 year olds. Thresholds were significantly reduced in the flanked-different condition, relative to flanked-same performance, for children in both the 3-6 year old (t_19_ = 2.648, p = 0.016, d = 0.59) and 7-12 year old groups (t_19_ = 2.987, p = 0.008, d = 0.67), as well as in adults (t_9_= 3.263, p = 0.010, d = 1.03), each with medium-to-large effect sizes. If we consider this reduction as a proportion of the elevation in crowding (i.e. the percentage reduction of the difference between unflanked and flanked-same conditions), then performance was improved by approximately 50% in both children and adults. Both children and adults thus show reduced crowding when flankers are reversed in their contrast polarity, demonstrating a common selectivity between the developing fovea, adult fovea, and adult periphery.

**Figure 5.**
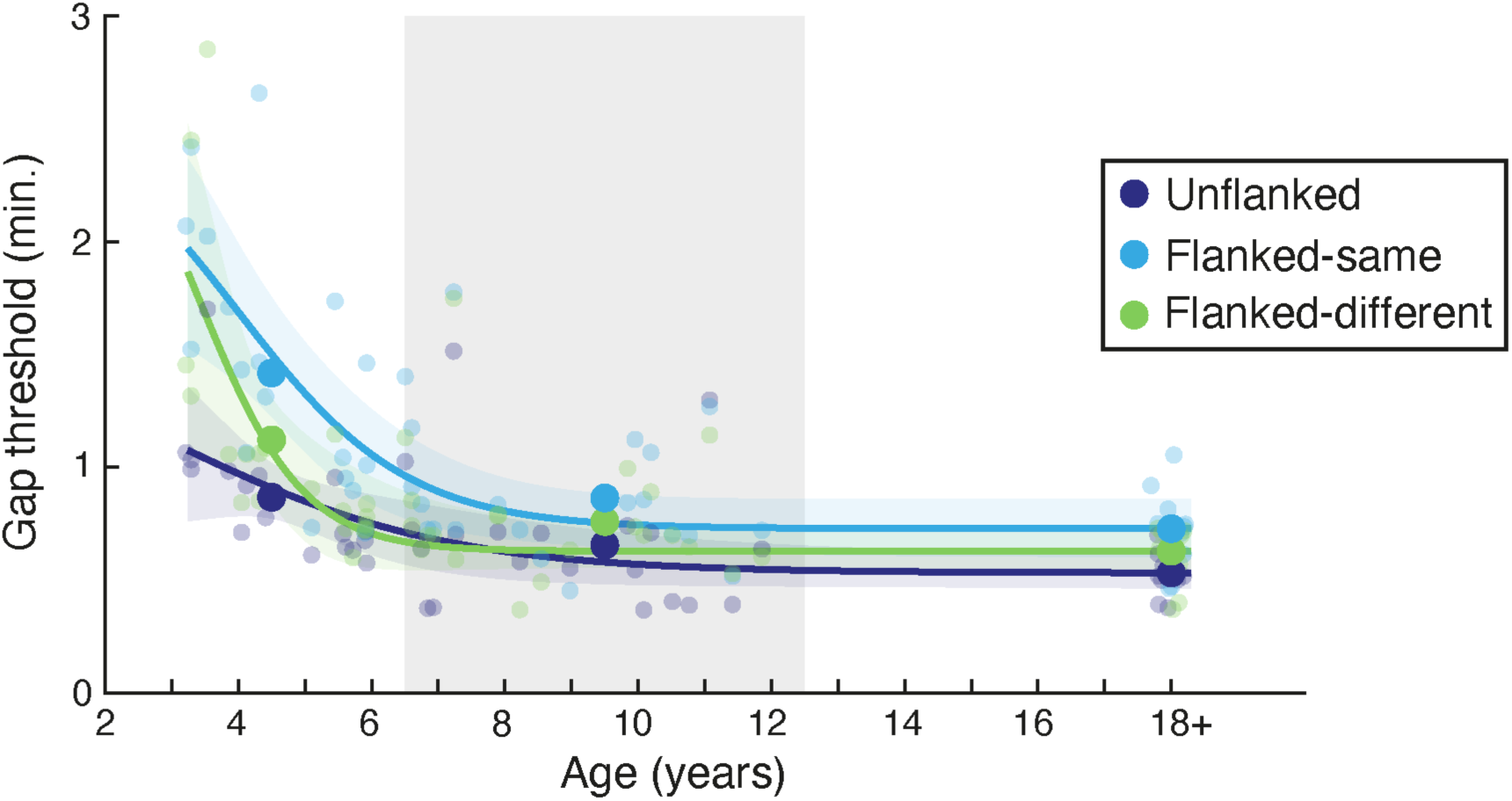
The release from crowding with age. Gap-size thresholds are in minutes of arc, plotted against age. Individuals are shown as small points and the mean within each age range (shown via grey/white bands) as large points, separately for the unflanked (dark blue), flanked-same (light blue), and flanked-different (green) conditions. Adult data has been collapsed to fall near 18 years. Lines show the best fitting logistic function to each condition, with shaded regions showing the 95% range of the fits to 1000 bootstrapped samples.

## Discussion

Our results confirm that foveal vision is disrupted by clutter in young children. Elevations in the spatial extent of crowding were most prominent in the youngest children – for 3-4 year old children, thresholds for the recognition of our VacMan stimuli were almost doubled in the presence of surrounding clutter. These elevations receded with age, but did not reach adult-like levels until the age of 7-8 years when clutter increased thresholds by a factor of about 1.4. This development is slower than that of acuity, which reached adult-like levels by around 5-6 years. The development of crowding was not however as slow as some estimates reported in prior studies. Despite these discrepancies in prior studies, our meta-analysis reveals good overlap in the combined developmental trajectory of 14 studies examining developmental crowding (13 prior studies and our own). This direct comparison was achieved after converting our thresholds to the metrics used by these studies, and gave an age at which crowding converged to adult-like levels around 7-8 years. Finally, we show that developmental elevations in foveal crowding also share a key property of peripheral crowding, with changes in target-flanker similarity (differences in contrast polarity) reducing crowding by 50% in all age groups. We conclude that developmental crowding derives from the same process as the disruptive effect evident in the adult visual system.

Our finding that foveal crowding becomes adult-like at 7-8 years is similar to estimates from several prior studies^21–23^, with our meta-analysis of 14 studies supporting this developmental timeline. Some discrepancies between individual studies in this meta-analysis were nonetheless apparent, most notably for one report that crowding remains elevated at 11+ years of age^20^. Our direct comparison of these measures across studies suggests that this discrepancy derives from an apparent underestimation of adult crowding levels. This underestimation could reflect differences in the procedures used to test adults vs. children, the exclusive use of psychophysically-trained adults as a comparison, or the use of relative measures to calculate the magnitude of crowding (whereby flanked and unflanked performance are combined to yield multiple or difference values). This latter issue was further evident in the conversion of our thresholds to the relative measures used in other studies (such as^15^), which led to more variable estimates than when acuity and crowding were independently measured, likely because the variance of these two components is combined (Figure S3). Studies using widely spaced elements^17,22^ also tended to underestimate crowding levels, particularly when target and flanker elements were dissimilar (bars surrounding letters)^17^ consistent with prior observations^44–46^. From this we recommend that developmental crowding is best measured against naïve adult observers, using tightly spaced stimuli, similar target and flanker elements, and with independent estimates of unflanked and flanked performance.

The studies compared in Figure 4 used a variety of stimuli and measurement procedures. Where these prior studies used multiple conditions to assess crowding, we selected those with the most standard configuration (typically 2-4 flankers surrounding a target). Arrays with larger numbers of elements have however been found to show a more pronounced elevation in the magnitude of foveal crowding for children than arrays with fewer elements^21,23^. Although it is possible that these additional elements could increase crowding effects, they may also introduce additional factors known to impair performance in cluttered arrays. For instance, rather than reflecting an immaturity in crowding itself, the increased complexity of these stimulus arrangements may increase uncertainty regarding the location of target stimuli^62,63^. This could in turn conflate the development of crowding with other processes such as spatial attention, which can interact with crowding^64^, though the two processes are clearly dissociable^65^. Other studies have reported slower developmental trajectories with children required to judge whether square-wave grating elements were horizontal or vertical^19^. Task comprehension with these more abstract stimuli may be an issue with children, and we suggest that the use of more gamified tasks is a useful means to clarify task requirements and ensure that children respond to the correct element.

Although it is clear that children make errors when judging the appearance of a fixated target in cluttered displays, it is not a given that these errors arise for the same reason as those found in adult peripheral vision. For instance, elevations in crowding could arise from changes in properties such as fixational stability^66^ smearing elements across the retina, a factor that contributes to adult foveal crowding levels^35,67^ and which becomes magnified in cases of infantile nystagmus^59,68,69^. Contrary to this possibility, here we observe that crowding is reduced in children when elements are dissimilar in their contrast polarity. This effect of target-flanker similarity matches prior observations of polarity-based reductions in the adult fovea^34,51,70^ and adult periphery^48–50^. This is also consistent with prior observations that children make the same kinds of errors in crowded arrays as adults in peripheral vision^41^, which can be accounted for using population pooling models of crowding^25,40^. In peripheral vision, this pooling process generates the ‘gist’ of the visual scene at the expense of fine detail^39,42^. We suggest it is the same pooling process that is being refined over this period in the fovea as the visual system seeks to balance the neural resources allocated to the recognition of fine detail vs. the extraction of summary statistics over a wider region to gain a simpler ‘gist’ of the scene.

Alternative accounts of crowding posit that these effects derive from grouping processes^37^, including variations in contrast polarity^70^. It is unclear what aspect of the visual system would be maturing by these accounts however, and given that these mechanisms are usually depicted as heavily top down^37^, one may expect a later maturation than that observed in the current study. Accordingly, processes that are often linked with grouping, such as the perception of illusory contours^71^ and the recognition of ambiguous two-tone images^72^, do not mature until around 10-12 years, suggesting that the grouping processes that modulate crowding should show a similarly late maturation. Attentional processes also appear to develop late, with abilities such as multiple object tracking^63^, attentional cueing^62^, and the visual search for feature conjunctions^73^ improving well into adolescence. This late maturation is similarly inconsistent with an attentional basis for crowding, as some have proposed^36^. These higher-level processes can nonetheless clearly interact with the magnitude of crowding, as above, and their role on developmental vision is important to ascertain.

Our results are consistent with a developmental cascade of maturation within the visual hierarchy^74,75^, whereby visual abilities that rely on higher levels of the cortical hierarchy tend to develop later^76,77^. Similar to prior estimates^12–14^, we find that acuity converges on adult-like levels at around the age of 5-6 years, earlier than the development of crowding. The developmental trajectory of acuity may then reflect the earlier maturation of retinal ganglion cells and primary visual cortex, given the link with acuity and these areas^78,79^. In contrast, the later maturation of crowding is consistent with observations that crowding relies on higher cortical levels than acuity, including candidate areas from V2-V4^80–83^. The exact properties that might be developing in these areas is less clear. Though crowding is often linked with receptive field size^81,82^, both pRF sizes and surround inhibition components in areas V1-V4 have been found to be adult-like by 6 years of age^84,85^. There are nonetheless some indications that cortical magnification around the fovea may develop later in childhood within areas V2 and V3^85^. The later development of these areas may then create the interplay between fine detail and ‘gist’ processing during development. As with arguments that the ‘gist’ of peripheral vision constitutes an efficient representation of complex visual scenes with limited neural resources^42,86,87^, it may be that the reduced acuity of infancy and early childhood creates a greater need for these summary representations, which then gives way to fine detail as the visual system matures. The slower reduction of this foveal ‘gist’ processing may then be driven by the later maturation of extrastriate cortical areas.

The slow maturation of crowding may also make this process more susceptible to disruptions during development. Similar proposals have been raised for motion perception, which develops late in childhood, with associated deficits in motion perception apparent in a range of developmental disorders^77,88^. Crowding is similarly elevated in a range of developmental disorders of vision including amblyopia^10,47^, nystagmus^59,68,69^, CRB1-related retinopathies^89^, and cerebral visual impairment^90^, which all have their onset in the age ranges where crowding is still maturing. Reports of elevated crowding in individuals with dyslexia^11,91,92^ and dyscalculia^93^ could similarly be the result of this late maturation, given their typical onset within this age range. Disruptions to the spatial and/or featural selectivity of the visual system would seem to create an increased drive for ‘gist’ processing in spatial vision. This in turn would limit processes that rely on the fine detail of the fovea, such as reading^8^. Indeed, increased inter-letter spacing has been found to reduce reading errors in both dyslexic^94,95^ and typically developing children^96^.

Altogether, our results suggest that the optimisation of spatial vision takes time – while acuity becomes adult-like around the age of 5-6, crowding continues to develop until 7-8 years of age. These developmental elevations in crowding share core patterns of selectivity with adult peripheral vision, suggesting that foveal vision faces similar trade-offs between the representation of fine vision and ‘gist’ during development. The slow development of this balance renders these abilities particularly susceptible to disruption and presents challenges for everyday tasks such as reading. Measurement of these processes is also a continuing challenge, in both typical and clinical cohorts. Here we suggest that crowding be measured in a child-friendly format with equivalent comparisons to adult data, using independent measures of stimulus conditions (e.g. unflanked and flanked performance), and with tightly spaced stimuli. The careful measurement of these effects will be important for future work attempting to examine whether other properties of crowding, such as its modulation by higher-level processes like grouping, develop at a similar rate.

## Acknowledgements

This work was funded by a UK MRC Career Development Award (MR/K024817/1) to JG and by a Wellcome Career Development Award (306332/Z/23/Z) to TD. Our thanks to Alexandra Kalpadakis-Smith for help with code development.

## Data availability statement

Data from the study is available at https://osf.io/d9fc8/ including code for analyses, which also require the Eccentric Vision toolbox, available from https://github.com/eccentricvision.

## Supplementary information for

### The influence of attentional lapses

Given suggestions that frequent attentional lapses can hinder the detection of visual abilities^97^, we examined the influence of these lapses on our data. We can quantify attentional lapses through the catch trials, whereby stimuli were presented at sizes three times larger than the QUEST estimate of threshold for that trial. Here we removed participants achieving less than 100% correct on these trials, independently for each of the unflanked and flanked-same conditions. There were not a great number of children who failed to meet this criteria in either case, though lapses were slightly higher in the flanked-same trials (as shown in Figure 3A). Amongst the 3-4 year olds, none were removed from the unflanked condition, with 2 removed from the flanked-same to leave 8. One 5-6 year old was removed from both conditions, leaving 21. One 7-8 year old was removed from the unflanked and two from the flanked-same, leaving 23 and 22, respectively. Of the 9-10 year olds, 3 were removed from the unflanked and 1 from the flanked-same, leaving 11 and 13, respectively. No 11-12 year olds were removed from the unflanked condition, and just one from the flanked-same to give 19 and 18, respectively. Six adults were removed from the unflanked condition to leave 24, and two from the flanked-same condition to give 28.

Thresholds from the remaining participants are shown in Figure S1. Mean values did not vary appreciably from those presented in the main analysis in Figure 3C. As before, isolated performance was significantly elevated relative to adult thresholds in the youngest children at 3-4 years (t_32_ = 4.853, p < 0.0001, d = 1.83), with no significant difference for any age groups beyond this (5-6 years: t_43_ = 1.495, p = 0.142, d = 0.45; 7-8 years: t_45_ = 1.575, p = 0.122, d = 0.46; 9-10 years: t_33_ = -0.872, p = 0.390, d = 0.32; 11-12 years: t_41_ = 1.239, p = 0.222, d = 0.38). The addition of flankers similarly produced clear elevations in threshold, which were significantly larger than adults at 3-4 years (t_34_ = 7.783, p < 0.0001, d = 3.12) and at 5-6 years (t_47_ = 3.141, p = 0.003, d = 0.91), but not at 7-8 years (t_48_ = 0.500, p= 0.619, d = 0.14) or beyond (9-10 years: t_39_ = 0.728, p = 0.471, d = 0.24; 11-12 years: t_44_ = 0.548, p= 0.586, d = 0.17). Given that the removal of participants with catch-trial performance below 100% does not alter the pattern of results, we conclude that differences in attentional lapse rates with age are unlikely to have played a major role in the developmental trajectories for acuity and crowding.

**Figure S1.**
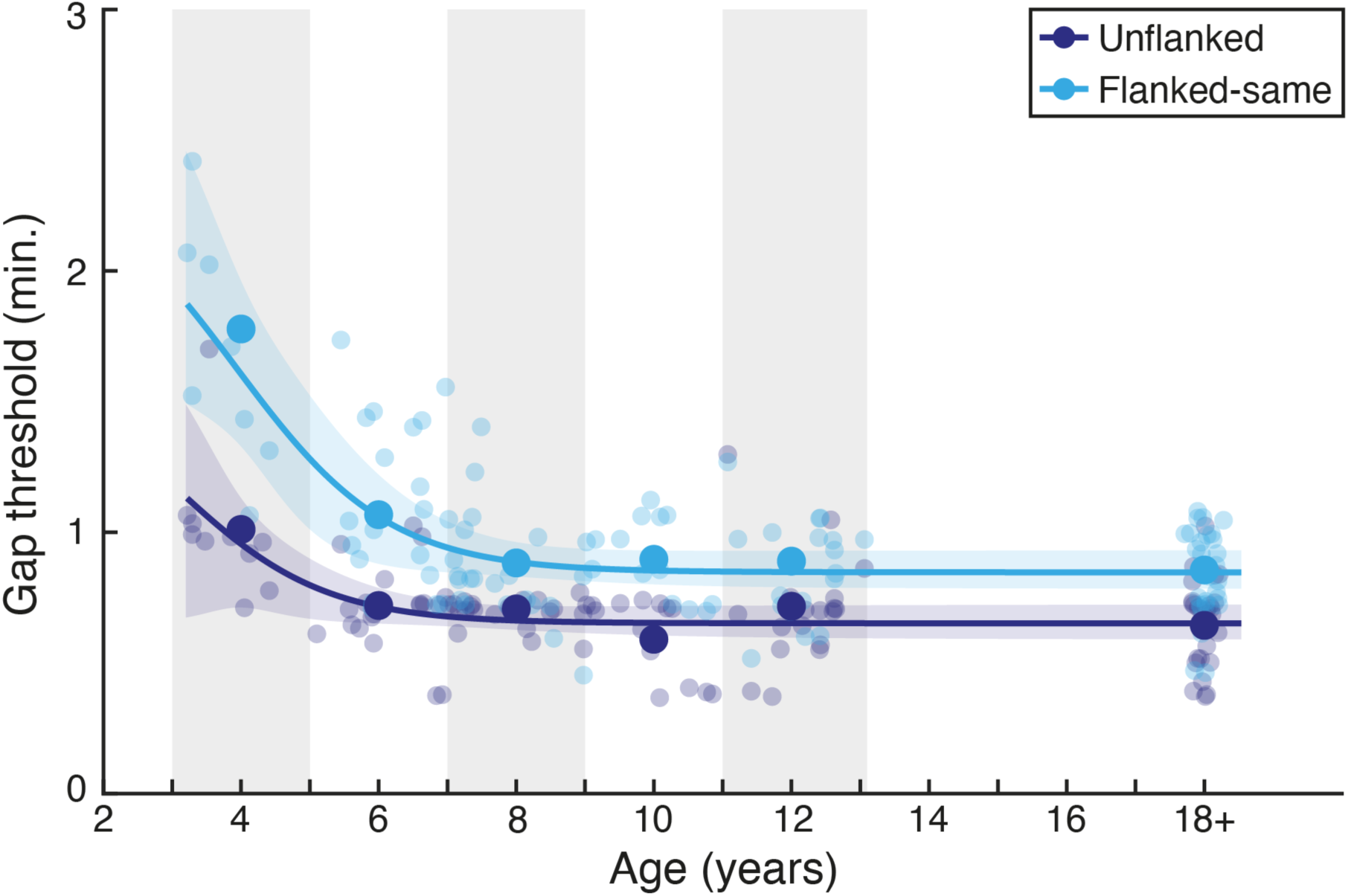
Gap-size thresholds plotted by age after filtering to remove individuals with performance on catch trials that was below 100% correct. As in Figure 3C, thresholds are plotted in minutes of arc, with individuals shown as small points and the mean in each age range (shown via grey/white bands) as large points, separately for the unflanked (dark blue) and flanked-same (light blue) conditions. Adult data has been collapsed to fall near 18 years. Lines show the best fitting logistic function to each dataset, with shaded regions showing the 95% range of fits to 1000 bootstrapped samples.

### Comparisons to prior studies using scaling vs. spacing approaches

Prior reports of the age at which crowding reaches adult levels, summarised in Figure 1, show an apparent divergence between studies where elements were scaled (as in the current study) and those using fixed stimulus sizes where inter-element separation was varied directly using elements of a fixed size (typically set as a multiple of unflanked acuity, using values measured prior to the measurement of crowding). Because two of our prior studies have taken these distinct approaches, we compared the resulting estimates of crowding to consider the influence of these measurement approaches.

The direct manipulation of inter-element spacing was previously used by Greenwood *et al*^47^ using the same VacMan stimuli to examine crowding in children aged 4-8. Using stimuli of a fixed size (2.5 times above acuity thresholds), Greenwood *et al*^47^ varied the centre-to-centre separation between elements to find the separation where performance reached 62.5% correct. Trial-by-trial data was extracted from this study and re-fit with psychometric functions for comparison to the current study. To consider the reliability of our estimates, we can further compare our values with a recent study^41^, where a scaling method similar to the current study was used with children aged 3-8 years. Because Kalpadakis-Smith *et al*^41^ used a higher threshold of 80% correct to measure the full extent of crowding, their trial-by-trial data was extracted and re-fit to obtain thresholds at 62.5% correct. Values from these two studies can then be compared with those of the current study by calculating the centre-to-centre separation of the elements in the current study at threshold. We can do so by multiplying our gap-size thresholds by 5 (to give stimulus diameter) and again by 1.1 (the scaled centre-to-centre separation) to give thresholds for the centre-to-centre separation between elements in minutes of arc.

The resulting individual values are plotted against those of the current study in Figure S2. Values from the current study again show a clear developmental trajectory, with thresholds from Greenwood *et al*^47^ and Kalpadakis-Smith *et al*^41^ largely interspersed amongst these values to follow this trajectory, aside from a small number of outliers. We conclude that thresholds obtained using a scaled approach (as in the current study and previously^41^) are comparable to those that vary inter-element separation directly^47^. If anything, the direct manipulation of inter-element separation would appear to produce more variable data, with two children from Greenwood *et al*^47^ in particular showing very large critical spacing estimates for their age. We suspect that this approach could lead to problems if acuity thresholds were over-estimated, which would make it difficult for children to reach ceiling performance in the flanked conditions (since stimuli may be too small to reach a high level of recognition, even at the largest separations). As the scaling approach measures flanked and unflanked performance independently, there are no such issues with the cross-dependency of thresholds.

**Figure S2.**
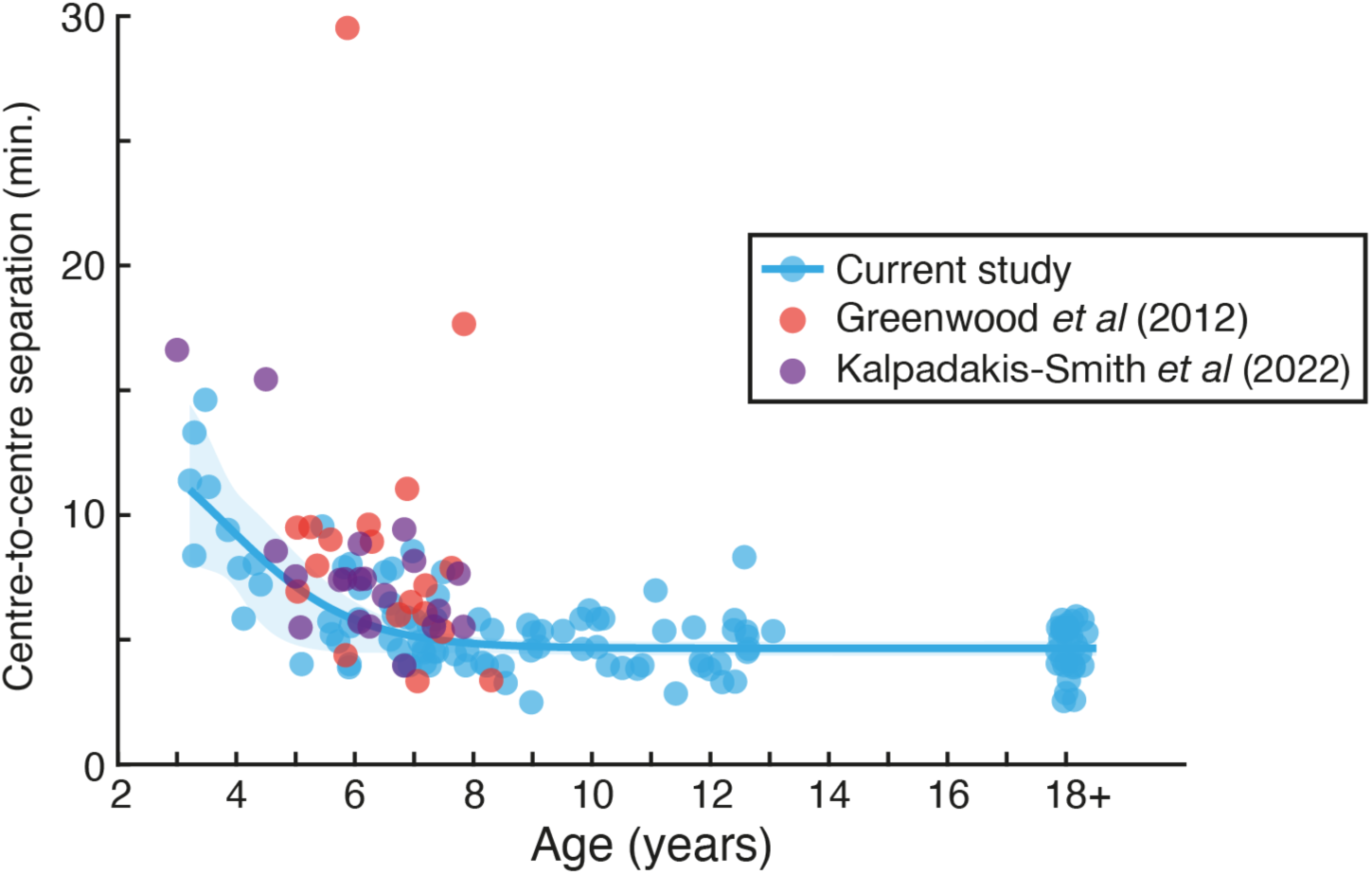
A comparison of different approaches to measure crowding. Thresholds from the flanked-same condition of the current study are shown in light blue, here expressed as the centre-to-centre separation between elements at threshold (in minutes of arc), along with the best-fitting logistic function for these values and a shaded region showing the 95% range of fits to 1000 bootstrapped samples. Thresholds are shown in equivalent centre-to-centre separation units from Greenwood *et al*^47^ in red (which measured inter-element separation directly) and Kalpadakis-Smith *et al*^41^ in purple (which used the same scaling approach as the current study). Each point represents an individual child.

### Meta-analysis of the developmental trajectory of crowding

To examine the correspondence between our estimates of the developmental trajectory of crowding and those of prior studies, we sought to compare these estimates directly. As well as the studies shown in Figure 1 (which only includes published studies with direct statistical comparison to adult data), here we also included studies without statistical comparison to adult values and unpublished work, allowing a broader view of the literature. Values from prior studies were extracted from the figures of each publication using the GRABIT tool for MATLAB. Mean values were taken in all but one case where individual values were plotted (as below). For comparison to our dataset, values were plotted either at the reported age values or at the middle of range values, where used. For each comparison, individual values from the present study were converted to the same units of measurement and re-fit with a logistic function. To compare values against those of the current study, we derived error terms for the logistic fit to our dataset (separately after conversion to each measurement unit) by taking the 95% range of values obtained by fitting the logistic function to 1000 bootstrapped samples of the data.

A common metric used in prior studies is to report crowding as multiples of acuity thresholds, measuring the elevation in crowding relative to baseline acuity levels. This approach was first taken by Atkinson *et al*^15^ using the Cambridge Crowding Cards (n=71 overall). Figure S3A shows that these values tend to be slightly lower than ours, consistent with the broader spacing of their elements (1.5× vs. 1.1× element size in the present study), which has been shown to underestimate the amount of crowding^44^. Thresholds nonetheless follow roughly the same developmental trend as the current study and fall within the 95% range of our fitted values for both sets of children. Similar measures were taken by Huurneman *et al*^61^ using acuity charts (n=75). Their most-closely spaced condition (1.0× element size) is also plotted in Figure S3A, which again follows largely the same developmental trend. Together, these estimates of crowded elevation are not widely divergent from those of the present study. Of note however is the individual variability in these estimates – whereas the individual values plotted separately for unflanked and flanked-same conditions clearly follow the developmental trajectory in Figure 3, their greater dispersal here makes the trajectory harder to discern. We suggest that it is better to take independent measures of performance and to compare them across conditions than to use estimates that combine multiple sources of error.

We can also compare our thresholds to prior work measuring crowding as the log difference between flanked and unflanked thresholds. Our data can be converted to this format by converting gap-size thresholds in minutes of arc to log units and then subtracting flanked thresholds from unflanked. Values from Norgett and Siderov^21^ obtained with a standard target-flanker arrangement (a letter target with 4 flankers, as in the current study, with n=89) are plotted in Figure S3B. These values lie close to ours, with the youngest children falling within our 95% range, though both older children and adults sit slightly below ours, likely driven by the wider spacing of 1.5× element size underestimating crowding levels, as above. Similar estimates can be seen in an earlier study using the same approach (n=103)^43^, also plotted in Figure S3B. Also included in this subplot is data from Facchin *et al* (n=252)^45^, who measured logMAR acuity using letter-chart arrangements at a range of inter-element spacings. For comparison, log thresholds were taken from the widest (2.0× element size) and narrowest (1.125×) centre-to-centre spacing conditions, which were then subtracted to give log difference values. These estimates again agree closely with our own.

**Figure S3.**
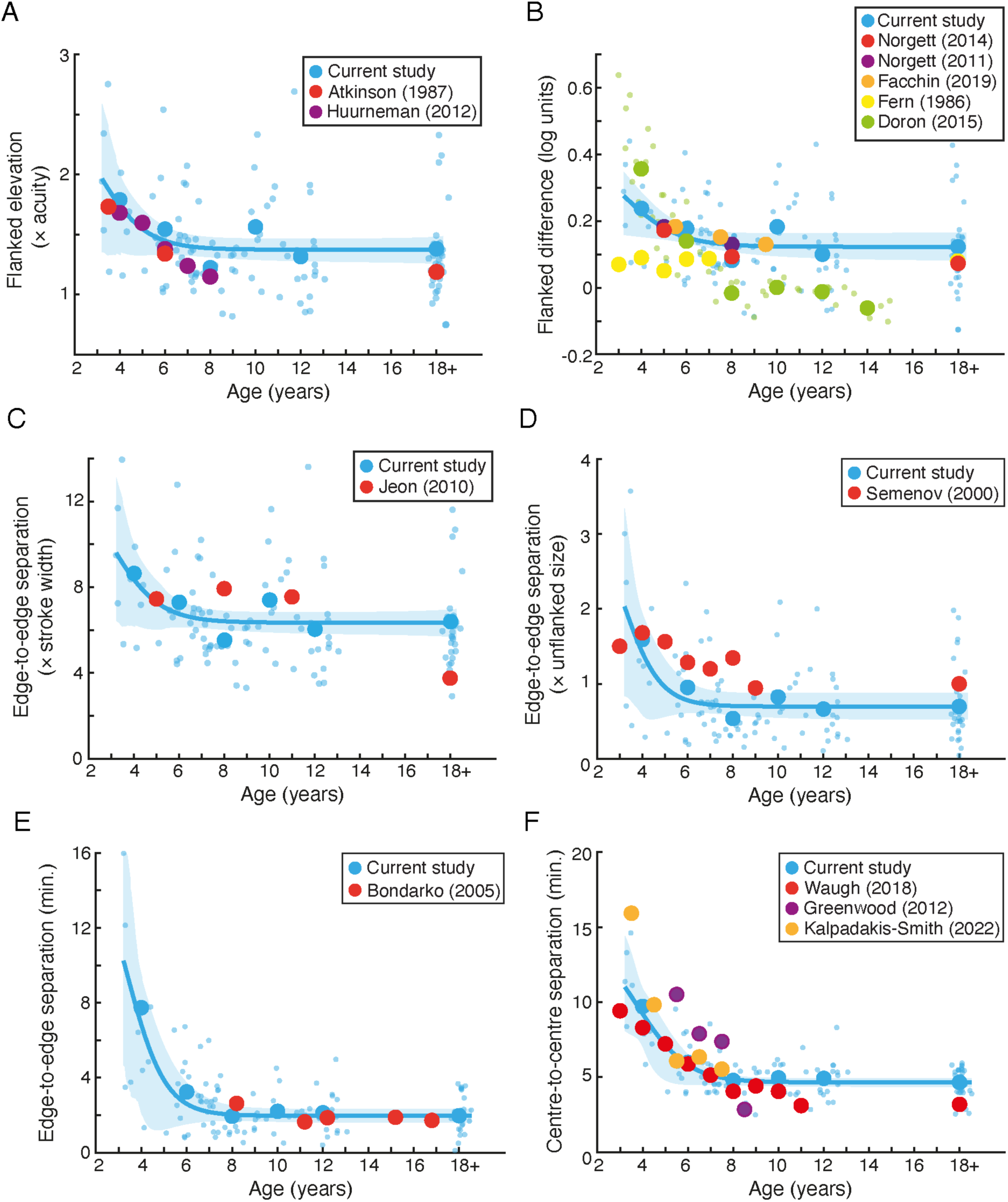
Comparison to prior datasets, plotted as a function of age on the x-axis. **A.** Crowding expressed in multiples of unflanked thresholds, with individuals shown as small blue dots and means for each age range (as in Figure 3) shown as large blue points. Comparison data is shown from Atkinson *et al*^15^ in red and Huurneman *et al*^61^ in purple. Lines show the best-fitting logistic function with shaded region indicating the 95% range of bootstrapped values. **B.** Crowding expressed as the difference between flanked and unflanked thresholds in log units, plotted against values from Norgett and Siderov^21^, Norgett and Siderov^43^, Facchin *et al*^45^, Fern *et al*^17^, and Doron *et al*^22^. **C.** Crowding expressed as the estimated edge-to-edge separation between elements, in multiples of stroke width, with comparison to data from Jeon *et al*^20^ in red. **D.** Crowding as edge-to-edge separation in multiples of element size, plotted against data from Semenov *et al*^18^. **E.** Crowding as edge-to-edge separation in minutes of arc, plotted against data from Bondarko and Semenov^19^. **F.** Crowding as the centre-to-centre separation between elements in minutes of arc, plotted against data from Waugh *et al*^23^, Greenwood *et al*^47^, and Kalpadakis-Smith *et al*^41^.

Differences emerge in the log difference values reported by Fern et al^17^, who used a C vs. O target discrimination task with letters either in isolation or flanked by four widely spaced flanker bars (at a centre-to-centre separation of approximately 1.5× target diameter, with n=121; Figure S3B). Although on its own their flanked data shows a clear developmental decline, their log difference values (with acuity subtracted) are uniformly low and largely unchanged with age. We suspect that this reflects an underestimation of crowding levels, for two reasons. First, the use of bars as flanker stimuli have been shown to underestimate crowding^21,44^, likely through the decreased target-flanker similarity of the bars to the target letters^30^. The large inter-element spacing would further decrease the level of crowding measured with these elements^44^. Differences are also evident in the values obtained by Doron *et al*^22^, measured with letter-chart arrays of Tumbling-E stimuli (n=46). Because individual values were plotted in this study, we recorded these and calculated mean crowding levels within each of the age ranges used in the current study, as plotted in Figure S3B. Although this approach gives an estimate for the age of maturity that is very similar to that of the present study, the estimates of crowding are higher than ours for the youngest children, likely due to the use of high numbers of elements causing uncertainty in the youngest children. An undershoot in crowding levels is also visible in the oldest children, likely driven by the letter-chart arrangement whereby each element is flanked by a maximum of 2 elements (depending on the position of the letter in each line). Although the absolute values diverge from ours in this instance, the developmental trajectory nonetheless remains largely the same.

The study with the oldest age of maturity for crowding comes from Jeon *et al*^20^, who took measures of the edge-to-edge separation between target and flanker elements, with elements presented at a size 1.2× acuity (measured initially) and varied in their separation (n=78). We can convert our values for comparison to theirs with some assumptions. The scaling of our elements meant that the edge-to-edge separation was always 0.1× the element size. We can however estimate what our values might have been had we varied spacing using fixed-size elements by taking the centre-to-centre separation between elements at threshold and subtracting the diameter of elements at the acuity threshold multiplied by 1.2, before dividing by the gap-size threshold to give units of stroke width. The resultant values are plotted against those of Jeon *et al*^20^ in Figure S3C. Only 1 of 3 estimates of childhood crowding coincide with our 95% range, though the largest discrepancy occurs in the adult data, where the estimates from Jeon *et al*^20^ are considerably lower than those of the adults measured with our approach. One possibility is that the adult sample collected by Jeon *et al*^20^ were more experienced with psychological testing than ours. Differences in the nature of the testing procedures used in children and adults could also give rise to this divergence. In comparison, our sample included predominantly adults who were inexperienced with psychological testing, with the same testing conditions used for both adults and children, and independent measures of the two abilities. The relative nature of these estimates (with flanked thresholds divided by unflanked) also makes it difficult to assess whether the difference reflects changes in crowding, acuity, or both.

Similar calculations allow comparison with the data of Semenov *et al*^18^, who varied the edge-to-edge separation between target and flanker bar elements presented at a size equal to acuity thresholds (n=141). Values were reported as edge-to-edge separation in multiples of unflanked whole-element size, which we can again calculate for our thresholds by taking the centre-to-centre separation between elements at threshold and subtracting the diameter of elements at the acuity threshold, before dividing by element diameter to give multiples of unflanked size. Semenov *et al*^18^ used a subjectively determined high level of performance to take threshold, which we attempted to match by taking thresholds at 90% correct (instead of 62.5% correct elsewhere). As shown in Figure S3D, the estimates of Semenov *et al*^18^ are elevated relative to ours in both children and adults, and show a period of continued elevation. Given the derivation of thresholds at high performance levels however, it is possible that some of this prolongation is due to attentional lapses, which are a primary cause of disrupted performance at the upper end of the psychometric function in particular^60,97^, which would also make the subjective estimation of asymptotic performance difficult.

Bondarko and Semenov^19^ conducted similar procedures to Semenov *et al*^18^, measuring the edge-to-edge separation between target and flanker elements, with elements presented at a size equal to acuity thresholds and varied in their separation (n=292). Here, values were reported as edge-to-edge separation in minutes of arc, which are more easily compared to ours (with the above assumptions regarding stimulus size), as plotted in Figure S3E. Here although their youngest 8-year-old participants show a slight elevation relative to ours, the bulk of the measurements align well with that of the current dataset. The age-range in our data nonetheless reveals that far greater changes in the developmental trajectory are evident before the age of 8 years. As with the estimates of Semenov *et al*^18^, the use of subjective fitting procedures to asymptotic data ranges may have led to the slight elevation in performance seen here with the 8 year old participants. Estimating a lower performance criterion may alter these estimates.

Finally, unpublished data from Waugh *et al*^23^ allows comparison to our dataset with fewer assumptions. Waugh *et al* used narrow number stimuli that were scaled using QUEST to obtain size thresholds. These estimates were converted to critical spacing values (n=241), similar to our computations shown in Figure S2. Taking their 3-element configuration (with one target and two flankers) as the closest to the majority of studies above, these estimates of the centre-to-centre separation between target and flanker elements at threshold can be compared to ours (Figure S3F). Both show a highly similar rate of decline in the size of these interference zones with age. We can further compare the data from Greenwood et al^47^ (n=19) and Kalpadakis-Smith et al^41^ (n=20) by binning these values into comparable age ranges and taking the means within each bin, also shown in Figure S3F. Values from Kalpadakis-Smith et al^41^ are again highly similar to both the current study and estimates from Waugh *et al*^23^, while those from Greenwood et al^47^ follow the same trend, though with increased variability, as noted above. The overall consistency of these developmental trajectories is nonetheless evident across these studies using similar approaches.

To combine these disparate metrics of crowding for meta-analysis, we used our dataset in each case as a reference. Each of the above 13 datasets were thus normalised by subtracting the minimum value from the mean values across age in our dataset (after conversion into the relevant metric), before dividing by the maximum mean value of our dataset. This set our dataset to span the range 0-1, with other datasets varying around this. The resultant values could then be combined in the same space. Mean values were taken within similar age ranges to those of the current study, which gave 19 estimates for 3-4 year-olds, 16 for 5-6 year-olds, 15 for 7-8 year-olds, 6 for 9-10 year-olds, 4 for 11-12 year-olds, 3 for 13-17 year-olds and 7 adult comparisons. Data for this meta-analysis are plotted in Figure 4 of the main text.

The analyses presented in Figure 4 of the main text include multiple contributions from individual studies in some of the age ranges. We also examined the pattern of data after this non-independence of individual datapoints was removed by averaging datapoints from individual studies within each of the above age ranges. This analysis restricted the contribution of each study to a maximum of 1 point per age group. Results from this analysis are plotted in Figure S4, which show a very similar developmental trend to that in Figure 4 of the main text. Crowding levels were again highest in the 3-4 year age range, which differed significantly from adult levels (t_16_= 4.423, p < 0.0001, d = 2.14). These levels remained significantly elevated at 5-6 years with a large effect size (t_15_ = 3.141, p= 0.007, d = 1.55), but dropped to levels that did not differ significantly from adults at 7-8 years (t_17_ = 1.339, p= 0.198, d = 0.64), and remained equivalent at 9-10 years (t_10_ = 0.676, p= 0.514, d = 0.40), 11-12 years (t_8_ = -0.601, p= 0.564, d = 0.41) and 13-17 years (t_6_ = -1.389, p= 0.207, d = 1.11). We conclude that the maturation of crowding at 7-8 years is a reliable feature of this meta-analytic dataset.

**Figure S4.**
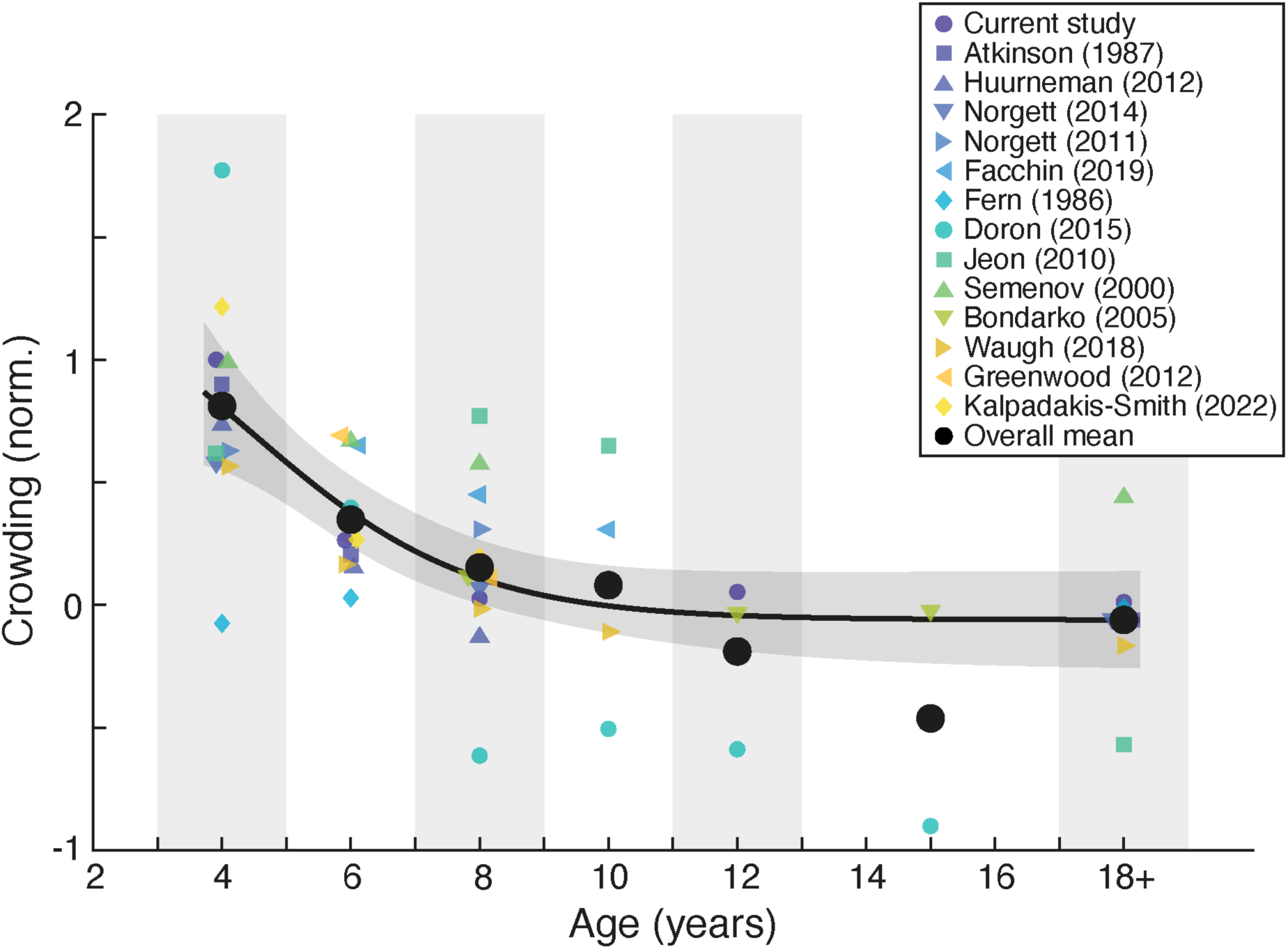
A meta-analysis of studies measuring the developmental trajectory of crowding. As in Figure 4 of the main text, crowding is plotted in normalised units, as a function of age on the x-axis. Small symbols show estimates from individual studies (here averaged within each age range prior to plotting), while the large black points show the mean within each age range (indicated via shaded regions). The black line is the best-fitting logistic function fit to all studies, with shaded region indicating the 95% range of the fits to 1000 bootstrapped samples. Some thresholds have been displaced on the x-axis for visibility.

